# SEPALLATA-driven MADS transcription factor tetramerization is required for inner whorl floral organ development

**DOI:** 10.1101/2023.05.23.541941

**Authors:** Veronique Hugouvieux, Romain Blanc-Mathieu, Michel Paul, Aline Janeau, Xiaocai Xu, Jeremy Lucas, Xuelei Lai, Antonin Galien, Wenhao Yan, Max Nanao, Kerstin Kaufmann, François Parcy, Chloe Zubieta

**Affiliations:** Laboratoire de Physiologie Cellulaire et Végétale, Université Grenoble-Alpes, CNRS, CEA, INRAE, IRIG-DBSCI, 17 rue des Martyrs, 38000 Grenoble, France; Plant Cell and Molecular Biology, Institute of Biology, Humboldt-Universität zu Berlin, Berlin, Germany; Current address: National Key Laboratory of Crop Genetic Improvement, Hubei Hongshan Laboratory, Huazhong Agricultural University, Wuhan, China; Current address: College of Plant Science and Technology, Wheat Genetics and regulomics, Huazhong Agricultural University, Wuhan, China; Structural Biology, European Synchrotron Radiation Facility, 71 Avenue des Martyrs, 38000 Grenoble, France

**Author notes:** **Material distribution:** The authors responsible for distribution of materials integral to the findings presented in this article in accordance with the policy described in the Instructions for Authors (https://academic.oup.com/plcell/pages/General-Instructions) are: Véronique Hugouvieux and Chloe Zubieta.

## Abstract

MADS genes encode transcription factors that act as master regulators of plant reproduction and flower development. The SEPALLATA (SEP) subfamily is required for the development of floral organs and plays roles in inflorescence architecture and development of the floral meristem. The SEPALLTAs act as organizers of MADS complexes, forming both heterodimers and heterotetramers *in vitro*. To date, the MADS complexes characterized in angiosperm floral organ development contain at least one SEPALLATA protein. Whether DNA-binding by SEPALLATA-containing dimeric MADS complexes are sufficient for launching floral organ identity programs, however, is not clear as only defects in floral meristem determinacy were observed in tetramerization impaired SEPALLATA mutants. Here, we used a combination of genome-wide binding studies, high resolution structural studies of the SEP3/AGAMOUS tetramerization domain, structure-based mutagenesis and complementation experiments in *sep1 sep2 sep3* and *sep1 sep2 sep3 ag-4* plants transformed with versions of *SEP3* encoding tetramerization mutants. We demonstrate that while SEP3 heterodimers are able to bind DNA both *in vitro* and *in vivo* and recognize the majority of SEP3 wild type binding sites genome-wide, tetramerization is not only required for floral meristem determinacy, but also absolutely required for floral organ identity in the second, third and fourth whorls.

## Introduction

MADS genes play central roles in the development of reproductive structures, from the specification of male and female cones in gymnosperms^1–5^ to the development of inflorescence architecture, ^6, 7^ determinacy of the floral meristem^8^ and the specification of floral organ identity in angiosperms^2, 9, 10^. The encoded MADS transcription factors (MTFs) bind to a highly conserved DNA sequence called a CArG box (CC-“Adenine-rich”-GG) as obligate dimers. The MTFs involved in reproductive development belong to the MADS type II, or MEF2 clade, and have a multidomain structure^11^. These domains consist of the highly conserved eukaryotic-specific DNA-binding MADS domain (M domain), a ∼30 amino acid alpha helical Intervening domain (I domain) critical for dimerization specificity^12^, a plant-specific coiled-coil Keratin-like oligomerization domain (K domain) and a largely unstructured and sequence-variable C-terminal domain (C domain). Based on this conserved domain structure, the type II MADS are also called MIKC and form two main groups which differ in their oligomerization capability. The “classic” MIKC^c^ group form dimers and tetramers and are important for reproductive structure development and organ identity. The MIKC* group, dimeric MTFs, has a more limited role in male gametophyte development^13, 14^. The well-established ABCE genetic model of floral organ identity requires the combinatorial activity of the MIKC^c^ genes^15^.

In floral organ development, the MADS genes are divided into the A (*SQUAMOSA, SQUA-like*), B (*DEFICIENS/GLOBOSA, DEF/GLO-like*), C (*AGAMOUS, AG-like*) and E (*AGL2/AGL6* or *SEP-like*) class genes^2^. The most recent common ancestor of seed plants likely contained both an A and E class ancestor which has been lost in gymnosperms^16^. Extant gymnosperms contain only B and C class MADS genes^2^, with B and C class-encoded MTFs directly interacting to specify the formation of male cones and the C-class MADS complexes specifying of female cones^4, 17^. While gymnosperm B and C-class MTFs are able to directly interact and likely form tetrameric complexes, this property has been lost in flowering plants, which require the angiosperm-specific SEPALLATA (SEP) subfamily (E class) to allow interaction of B and C MTFs for third whorl organ specification (stamen). Likewise, female organ development (carpel) in the fourth whorl of angiosperms also requires the E class SEP subfamily in addition to the C class MADS^5, 18, 19^. The identity programs for the perianth organs (sepals and petals) in angiosperms require an A class MADS gene, with sepal formation in *Arabidopsis* dependent on A and E class MTFs and A, B and E class MTFs required for the determination of second whorl petal identity^15, 20^. The tetramerization domain was recruited early in seed plant evolution, with more promiscuous tetramerization putatively occurring between different MIKC^c^ MTFs in ancestral species. Loss of direct tetramerization capability between B and C class MTFs occurred after the gymnosperm-angiosperm split, with the E class SEPALLATAs taking over the tetramerization function. Based on these data, tetramer formation has been long hypothesized to be key for reproductive organ development triggered by MTFs. However, direct evidence for this has remained elusive, due in part to the limitations in protein-protein interaction studies which mainly identify binary interactions, the difficulty in characterizing transcriptionally active MADS complexes *in vitro* and *in vivo* and the study of loss-of-function mutants which are not sufficient to probe development as a function of different oligomerization states.

In *Arabidopsis thaliana*, which contains four *SEP* genes, triple (*sep1 sep2 sep3*) and quadruple (*sep1 sep2 sep3 sep4*) mutants display strong floral phenotypes, including loss of meristem determinacy and homeotic conversion of floral organs into sepaloid or leaf-like organs, respectively ^21–23^. Furthermore, *35S*-driven expression of B and/or C class MADS genes alone is not sufficient to launch floral organ identity programs and the concurrent expression of a *SEP* gene is required for the formation of ectopic floral organs^21, 22, 24, 25^. At the molecular level, this suggests that SEP-containing heterodimers or tetramers are required for proper MADS function. Extensive yeast 2-hybrid experiments have demonstrated that SEP MTFs are able to oligomerize with class A, B and C MTFs. Yeast 3-hybrid and *in vitro* experiments further demonstrate the formation of SEP-containing heterotetrametric complexes, 2-site DNA binding and DNA-looping^26–29^. Recent studies using a tetramerization-impaired *SEP3* allele, *SEP3^Δtet^*, expressed in the *sep1 sep2 sep3* mutant background attempted to decouple DNA binding and oligomerization state. This work demonstrated that robust tetramerization is required for floral meristem determinacy, but left open the question as to the role of tetramerization in floral organ identity as second and third whorl organ identity programs were not affected and fourth whorl organ identity was only partially perturbed compared to the loss-of-function *sep1 sep2 sep3* triple mutant^30^. Examination of genome-wide binding using sequential DNA affinity purification and sequencing (seq-DAP-seq) indicated two-site co-operative binding at certain loci by SEP3/AG tetramers, the complex required for fourth whorl organ identity, which was lost in complexes containing SEP3^Δtet^, further suggesting that hetero-dimerization of E and C class MTFs may be sufficient for carpel identity^30^.

In order to address the fundamental question of the physiological role of tetramerization in flower development, we performed structural, biochemical, and *in vivo* experiments to correlate oligomerization state with DNA-binding and physiological function. Using structure-based design, we generated SEP3 and AG mutants with strongly abrogated tetramerization capability and compared their DNA-binding and ability to rescue the *sep1 sep2 sep3* triple mutant phenotype. These results demonstrate that while SEP3-containing dimeric complexes bind many of the same sites as SEP3-containing tetramers genome-wide, they are unable to restore organ identity in the second, third and fourth whorls. Short-range binding site co-operativity based on intersite spacing enrichment is strongly reduced in the tetramerization mutants in seq-DAP-seq experiments and band-shift assays, pointing to a mechanism of DNA looping as important for proper gene regulation in organ identity. Taken together, these data show the absolute requirement of tetramerization for organ specification and proper cellular identity of petals, stamen and carpels.

## RESULTS

### SEP3 oligomerization, structural studies and mutant design

The importance of MADS tetramerization in floral organ development has been extensively investigated, most recently in the context of the central role of *SEP3*, the only *SEP* gene able to fully complement organ identity as a single allele in the *sep1 sep2 sep3* triple mutant (Supplemental Figure S1). Using a natural splice variant impaired in tetramerization, SEP3^Δtet^, *in vitro* DNA-binding studies demonstrated the loss of co-operative two-site DNA binding for the SEP3/AG heterocomplex, responsible for fourth whorl development and determinacy^31^. Relatively mild effects were observed *in vivo* in complementation assays, with phenotypes restricted to the fourth whorl and indeterminacy of the floral meristem^30^. Genome-wide binding studies using ChIP-seq (Supplemental Figure S2A) coupled with comparative RNA-seq of *SEP3* and *SEP3^Δtet^*expressing plants (Supplemental Figure S2B and C) were consistent with the observed phenotypes and highlighted relatively few differences in DNA-binding or gene regulation between *SEP3* and *SEP3^Δtet^ in vivo*. This may be due to the residual ability of *SEP3^Δtet^* to tetramerize *in vivo* with MADS partners, and would account for the rescue of second and third whorl organ identity as well as the partial restoration of fourth whorl identity as previously described^30^.

In order to better design SEP3 mutants no longer able to tetramerize, we solved the structure of the physiologically relevant MADS heterotetrameric SEP3/AG K domain complex, using seleno-methionine derivatized protein and single anomalous dispersion (SAD) phasing. A partial structure was autobuilt using ARP/wARP^32^ and subsequently used for molecular replacement of a higher resolution native SEP3/AG dataset. The protein complex crystallized in spacegroup C222_1_ with 8 molecules per asymmetric unit. The resolution was 2.4Å for the native dataset and the refined model exhibited very good geometry and no residues in disallowed regions of the Ramachandran plot (Table 1). As shown in Figure 1A, the crystal structure of SEP3/AG contains the complete K domains of SEP3 and AG, a small portion of the I domain and several residues of the C domain, with the tetramer adopting a cross-like configuration with outstretched alpha helical “arms”. The overall structure is very similar to the previously described SEP3 homotetramer (PDB 40XO), however the tetramer of SEP3/AG exhibits additional salt bridge interactions along the protein-protein interface (Figure 1B)^29^. Structural comparisons between SEP3/AG and SEP3 tetramers (Figure 1C) reveals a slight change in orientation of the alpha helical arms, with SEP3/AG exhibiting a more planar orientation of the N-terminal helices.

**Figure 1.**
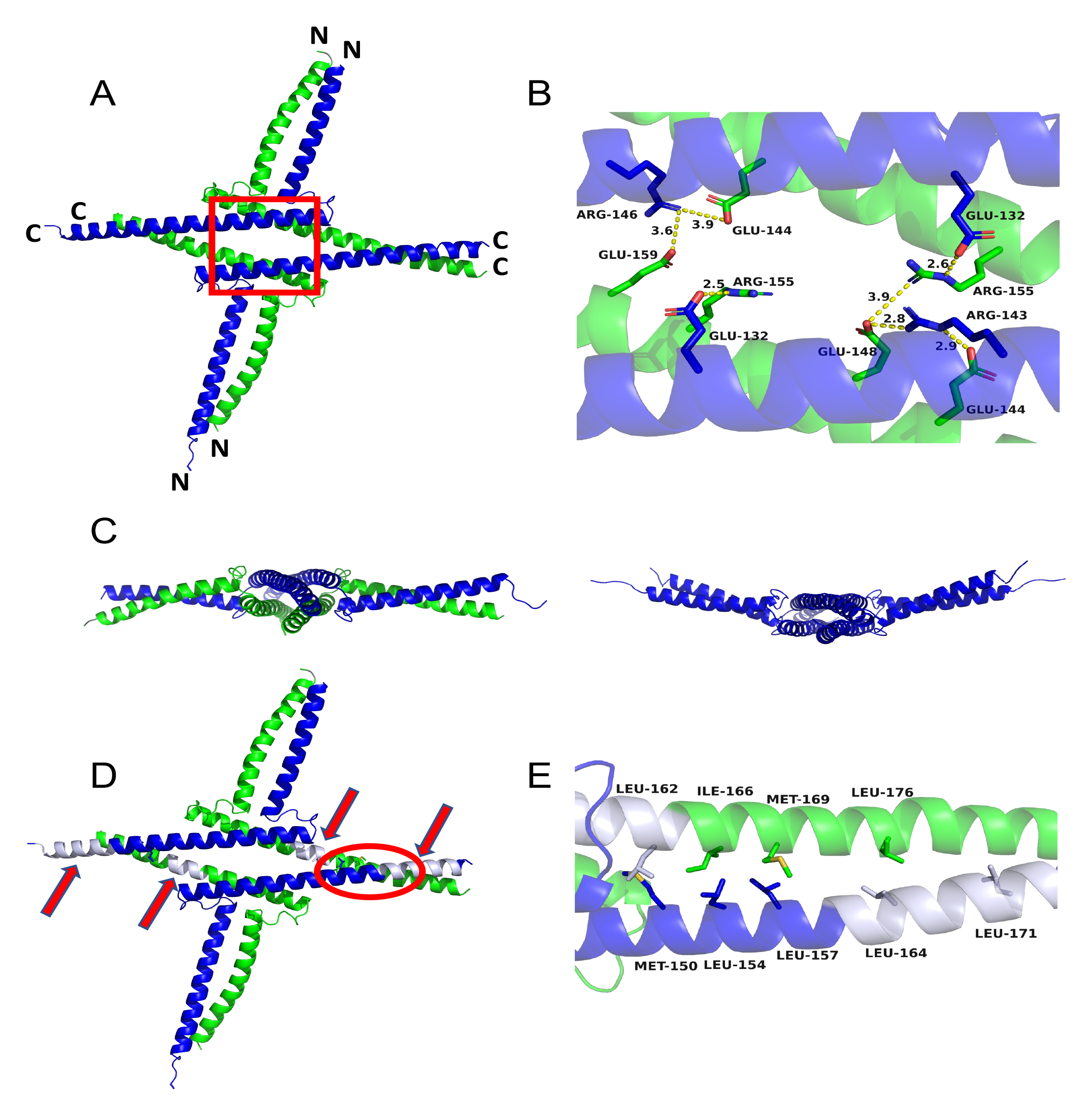
Structure of SEP3/AG heterotetramer. **A.** SEP3 (blue) and AG (green) tetramer shown as a cartoon. N-and C-termini are labeled. The red box denotes the zoomed in region in **B**. **B.** Close-up of salt bridges at the tetramerization interface of SEP3 and AG. Residues are labeled and salt bridges are shown as dashed yellow lines with distances shown. **C.** View of SEP3/AG (left) and SEP3 (right) tetramers looking down the C-terminal alpha helices. SEP3 homo-tetramer exhibits a curvature as compared to SEP3/AG heterotetramer. **D.** Cartoon representation as in **A**, with the deletion mutations SEP3^Δtet^ and AG^Δtet^ colored in gray and indicated by red arrows. The circled region is rotated for clarity and shown in **E**. **E.** Close-up view of the hydrophobic tetramerization interface between the C-terminal alpha helices of SEP3 and AG. Hydrophobic residues are labeled. The SEP3^Δtet3M^ mutations target the leucine zipper, with M150A, L154A and L157A and a deletion of residues 161-174 all affecting the tetramerization interface.

**Table 1.** Data collection and refinement statistics

|  | SEP3-AG |
| --- | --- |
| **Data collection** |  |
| Space group | C222<sub>1</sub> |
| Cell dimensions |  |
| *a*, *b*, *c* (Å) | 101.3, 138.4, 180.2 |
| $\alpha$, $\beta$, $\gamma$ (°) | 90, 90, 90 |
| Resolution (Å) | 82.-2.4 (2.45-2.40)* |
| *R*<sub>sym</sub> or *R*<sub>merge</sub> (%) | 9.7 (163) |
| *I* / σ*I* | 11.3 (0.9) |
| Completeness (%) | 99.3 (93.2) |
| Redundancy | 3.8 (3.7) |
| CC(1/2) | 99.9 (34.7) |
| **Refinement** |  |
| Resolution (Å) | 20.-2.4 (2.45-2.4) |
| No. reflections | 46989 |
| *R*<sub>work</sub> / *R*<sub>free</sub> | 23.6/27.9 (33/35) |
| No. atoms | 6175 |
| Protein | 6160 |
| Water | 115 |
| Other ligands | - |
| *B*-factors |  |
| Protein | 82 |
| Water | 66 |
| Other ligands | - |
| R.m.s. deviations |  |
| Bond lengths (Å) | 0.007 |
| Bond angles (°) | 1.48 |

\* refers to the highest resolution shell

Examination of the highly conserved hydrophobic leucine zipper critical for tetramerization allowed us to design point mutations to generally target the tetramerization interface for SEP3-containing MADS complexes (Figure 1D and E). In addition to the SEP3^Δtet^ mutation that deletes residues 161-174, three additional point mutations were introduced to create a new mutant, SEP3^Δtet3M^, carrying the 161-174 amino acid deletion and mutations M150A, L154A and L157A (Figure 1D and E). The introduced mutations are all present at the predicted protein-protein interface for the heterotetrametric MADS protein complexes, suggesting that these mutations should universally disrupt SEP3-dependent tetramerization. In addition, the *ag-4* allele, which encodes a version of AG lacking residues 159-172, which we refer to as AG^Δtet^, was mapped to the SEP3/AG structure (Figure 1D and E). This deletion mutant affects the N-terminal portion of the SEP3/AG tetramerization interface. Based on the structure of SEP3/AG, the combination of SEP3^Δtet^ and AG^Δtet^ results in a SEP3/AG complex unable to tetramerize as it completely lacks the interface required for stable tetramer formation.

### *In vitro* characterization of SEP3^Δtet^ and SEP3^Δtet3M^ protein complexes

Electrophoretic mobility shift assays (EMSA) were performed to evaluate the impact of mutations on MADS complex formation and DNA binding. First, we tested different SEP3 mutants with AG and AG^Δtet^. As shown in Figure 2A, EMSAs performed with SEP3**^Δ^**^tet3M^/AG or with SEP3**^Δ^**^tet^/AG^Δtet^ confirmed that the mutations completely abolish heterotetramer formation *in vitro*, as observed by the complete disappearance of the band corresponding to tetrameric complexes. As previously reported, SEP3**^Δ^**^tet^ was only partially impaired in its ability to form heterotetramers with AG^30^. In all cases, mutations did not impair dimer formation and binding to DNA as confirmed by the presence of a strong band corresponding to migration of a dimer bound to DNA.

**Figure 2.**
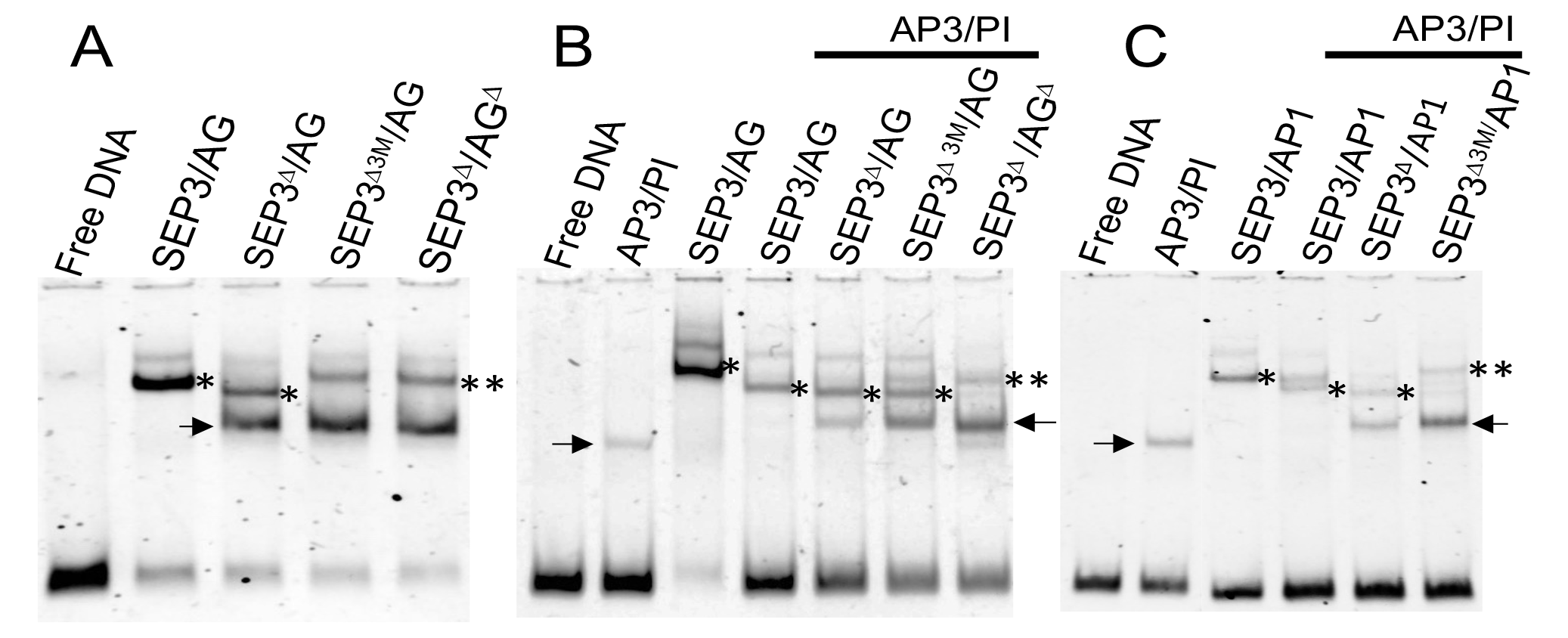
Electrophoretic mobility shift assays for MADS complexes using DNA with two CArG-box MADS binding sites. **A.** Fourth whorl C and E MADS complexes are shown with different SEP3/AG complexes forming dimers and tetramers. The wild-type SEP3/AG complex binds DNA as a tetramer, SEP3^Δtet^/AG binds as a mixture of dimeric and tetrameric species. SEP3^Δtet3M^/AG and SEP3^Δtet^/AG^Δtet^ bind DNA as one or two dimers. **B.** MADS B+C+E complexes important for third whorl organ identity with SEP3/AG shown for comparison. The mixture of SEP3^Δtet3M^/AG/AP3/PI shows a reduction in tetramer formation while the mixture of SEP3^Δtet^/AG^Δtet^/AP3/PI does not bind DNA as a tetramer but as one or two dimers. **C.** MADS A+B+E complexes important for second whorl organ identity. The mixture of SEP3^Δtet3M^/AP1/AP3/PI shows a reduction in tetramer formation as compared to SEP3/AP1/AP3/PI or SEP3^Δtet^/AP1/AP3/PI. The AP3/PI heterodimer and SEP3/AP1 hetero-tetramer are shown for comparison. SEP3^Δtet^ is denoted as SEP3^Δ^ and AG^Δtet^ as AG^Δ^ for simplicity in the figure. Arrows indicate dimeric complexes, * indicates a tetramer and ** indicates two dimers.

Next, we evaluated SEP3 and AG mutants for their ability to affect complexes important for third whorl (AP3/PI/AG/SEP3) organ identity and SEP3 mutants for second whorl (AP1/AP3/PI/SEP3) identity. When co-expressed with AG/AP3/PI (Figure 2B) or AP1/AP3/PI (Figure 2C), SEP3**^Δ^**^tet3M^ was more strongly affected in tetramer formation than SEP3**^Δ^**^tet^, as indicated by a less intense upper tetramerization band and a more intense lower band corresponding to a dimer-bound DNA complex. Interestingly, heterocomplex formation was completely abolished between AP3/PI/AG^Δtet^/SEP3**^Δ^**^tet^, as no band corresponding to the hetero-complex was observed (Figure 2B). Taken together, the data show that the SEP3**^Δ^**^tet3M^ mutant, or the combination of SEP3**^Δ^**^tet^ and AG^Δtet^, provoke much stronger tetramerization defects *in vitro* than SEP3**^Δ^**^tet^ alone, as effects on co-operative DNA-binding were observable for MADS complexes involved in second, third and fourth whorl, and third and fourth organ identity, respectively.

### Comparison of DNA-binding by SEP3^Δtet3M^/AG and SEP3^Δtet^/AG^Δtet^ complexes

Based on these results, SEP3^Δtet3M^ containing complexes and the SEP3^Δtet^/AG^Δtet^ complex almost completely abolish tetramer formation as compared to SEP3^Δtet^ on individual binding sites. We used seq-DAP-seq for a genome-wide comparison of the binding of the three complexes (SEP3^Δtet^/AG, SEP3^Δtet3M^/AG and SEP3^Δtet^/AG^Δtet^) at regions bound by the wild type SEP3/AG. Our previous analysis using seq-DAP-seq showed that the SEP3^Δtet^ mutation reduced both the binding affinity and the preference for specific CArG-box spacing (36, 46, 56 bp)^31^. We considered a region to be ‘unbound’ by a mutant complex for which the binding intensity is decreased by at least a factor two (Coverage Fold Reduction CFR > 2) as compared to SEP3/AG binding (Figure 3). Most regions (> 4672, 74%) were bound with similar intensity (CFR relative to SEP3/AG < 2) by all complexes (Figure 3A). The presence of SEP3^Δtet3M^ or AG^Δtet^ in the heterocomplex led to an additional binding reduction as compared to the SEP3^Δtet^ mutation alone with 690 and 686 regions bound by SEP3^Δtet^/AG and not by SEP3^Δtet3M^/AG or SEP3^Δtet^/AG^Δtet^, respectively. Of these newly lost regions, 377 were shared by SEP3^Δtet3M^/AG and SEP3^Δtet^/AG^Δtet^ complexes. Moreover, genome-wide, the median binding intensity of SEP3^Δtet3M^/AG and SEP3^Δtet^/AG^Δtet^ relative to that of the SEP3^Δtet^/AG mutant had a stronger decrease at regions containing a preferred CArG-box intersite spacing as compared to regions with no preferred CArG-box intersite spacing (Wilcoxon test, *P*=7×10^-14^ and 0.0005 for SEP3^Δtet3M^/AG and SEP3^Δtet^/AG^Δtet^, respectively), suggesting that SEP3^Δtet3M^/AG and SEP3^Δtet^/AG^Δtet^ are less able to bind interspaced CArG-boxes as compared to SEP3^Δtet^/AG (Figure 3B). The list of genes associated with at least a two-fold reduction in binding between the wild type and tetramerization mutants is given in Table SI and includes genes such as *KANADI2 (KAN2)*, which encodes a TF involved in carpel and ovule development and the establishment of polarity of floral organs^33, 34^, *JAGGED (JAG)*, which encodes a zinc-finger TF important for stamen and carpel development^35, 36^ and *INNER NO OUTER (INO*), a gene encoding a YABBY TF implicated in ovule integument development^37^. Taken together, these *in vitro* genome-wide binding comparisons demonstrate a small but statistically significant impairment of SEP3^Δtet3M^/AG and SEP3^Δtet^/AG^Δtet^ DNA-binding compared to SEP3^Δtet^/AG, and a strong decrease in regions bound by tetrameric SEP3/AG complexes. This suggests a genome-wide quantitative relationship between tetramer formation and access to regions showing specific intersite spacing at certain loci, with these regions putatively acting as important organ identity determinants.

**Figure 3.**
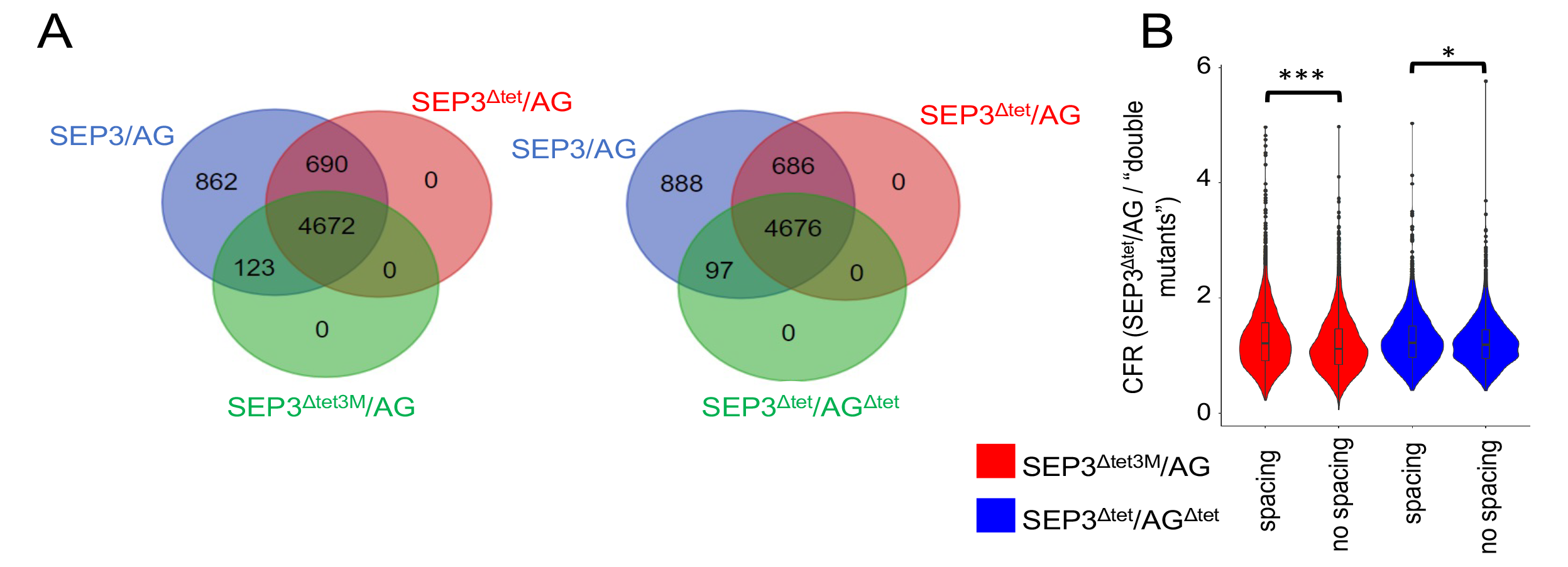
Genome-wide DNA binding comparisons determined by seq-DAP-seq for SEP3/AG wild-type and mutant complexes. **A**. Venn diagrams showing regions specifically bound by SEP3/AG (blue), SEP3^Δtet^/AG (red) and SEP3^Δtet3M^/AG (green; diagram on the left) or specifically bound by SEP3/AG (blue), SEP3^Δtet^/AG (red) and SEP3^Δtet^/AG^Δtet^ (green; diagram on the right) complexes. Regions specifically bound are defined as having a binding intensity at least twice greater for a complex relative to the other complexes. **B**. Binding intensity ratio of SEP3^Δtet^/AG to SEP3^Δtet3M^/AG (red) and SEP3^Δtet^/AG to SEP3^Δtet^/AG^Δtet^ (blue) over the 6,347 regions bound by SEP3/AG. The change in binding intensity is more significant for regions with a specific CArG-box intersite spacing (*n*=2270) than for region with no spacing (*n*=4077) for both SEP3^Δtet3M^/AG and SEP3^Δtet^/AG^Δtet^ versus SEP3^Δtet^/AG (Wilcoxon test, ***: P <10^-5^, *: P <10^-3^).

### Impact of MADS tetramerization mutants in floral organ development and cell identity

Based on *in vitro* data, the series of mutations targeting the tetramerization interface were used to assess the importance of MADS tetramerization in the different floral organ development programs. We generated *sep1 sep2 sep3* plants expressing *SEP3*, *SEP3****^Δ^****^tet^* or *SEP****^Δ^****^tet3M^*, and *sep1 sep2 sep3 ag-4* plants expressing *SEP3****^Δ^****^tet^* and analyzed the overall morphology of each floral organ and the surface cell identity in the second, third and fourth whorls by scanning electron microscopy (SEM) (Figures 4 and 5). The *sep1 sep2 sep3* mutant exhibited conversion of all floral organs to sepaloid structures that showed numerous stomata and typical elongated cells at their surfaces (Figures 4 and 5, first columns). The lack of determinacy of the floral meristem results in the continuous generation of a new “flower” made of sepaloid organs in the fourth whorl (Figure 4)^21, 30^. Flowers of *sep1 sep2 sep3* plants expressing *SEP3* were fully complemented (Figure 4, second column) and showed WT petals, stamens and carpels in whorls 2, 3 and 4, respectively, with conical cells, pollen grains and stigmatic papilla, style and replum cells at the appropriate organ surface (Figure 5, second column). As previously described, *SEP3****^Δ^****^tet^* expression in *sep1 sep2 sep3* was able to fully complement petal and stamen formation in whorls 2 and 3, but only partially complemented whorl 4, which exhibited two unfused carpel-like structures and indeterminacy (Figures 4 and 5, third column)^30^. In contrast, flowers of plants expressing *SEP3****^Δ^****^tet3M^* (Figures 4 and 5, fourth column) showed significant defects in whorls 2 and 3 compared to *SEP3****^Δ^****^tet^* expressing plants, and no carpel-like structures in whorl 4. In the second whorl, the petaloid organs were much shorter than WT petals and remained green (Figure 4). No stomata cells were visible and conical cells were only occasionally observed by SEM (Figure 5). In the third whorl, only immature greenish stamen could be observed (Figure 4). Small blisters at the organ margin that resemble developing pollen sacs were also noted but no pollen grains were produced (Figure 5). The number of organs in whorls 2 and 3 was not affected in these plants, with four and six organs in the second and third whorls, respectively. These data show that reducing the ability of SEP3 to tetramerize results in increasingly strong defects in floral organs, including incomplete organ differentiation and cell identity, notably in whorls 2 and 3 that were unaffected in the *SEP3****^Δ^****^tet^* expressing plants.

**Figure 4.**
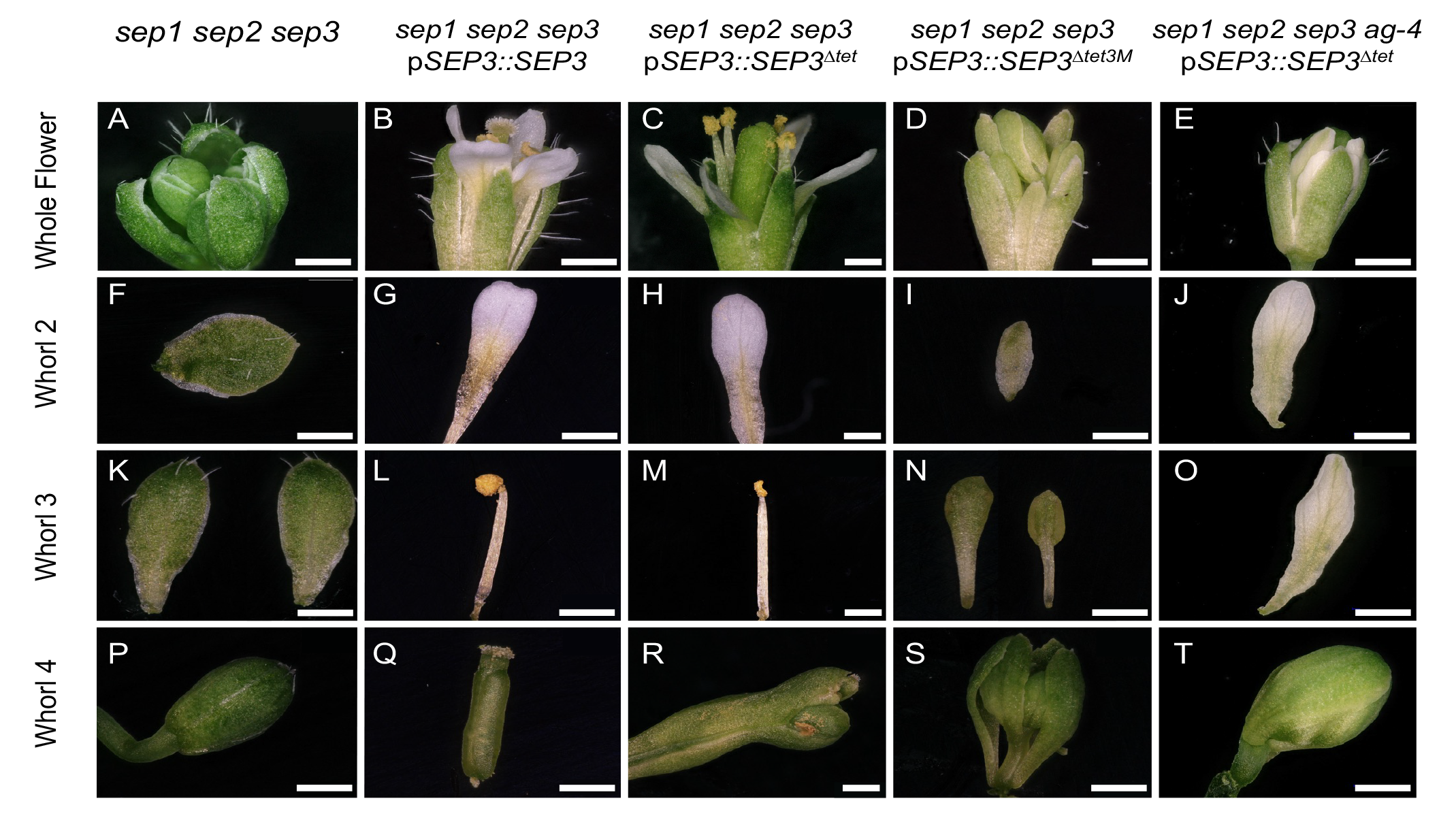
The flower and second, third and fourth whorl floral organs in *Arabidopsis* expressing wild-type and MADS mutants. **A-E.** Representative whole flowers in, from left to right, *sep1 sep2 sep3*, *sep1 sep2 sep3* expressing *SEP3, SEP3^Δtet^* or *SEP3^Δtet3M^* and *sep1 sep2 sep3 ag-4* expressing *SEP3^Δtet^* as labeled (top). Representative organs of whorl two **(F-J)**, whorl three **(K-O)** and whorl four **(P-T)** for each genotype described above. *sep1 sep2 sep3* expressing *SEP3* plants are fully complemented and show WT organs*. SEP3^Δtet3M^* expressing plants exhibit strong floral organ phenotypes in the second, third and fourth whorls with immature green organs. The combination of *ag-4* and *SEP3^Δtet^* triggers the complete transformation of stamen into petals in the third whorl and indeterminacy in the fourth whorl. Scale bars indicate 500 µm.

**Figure 5.**
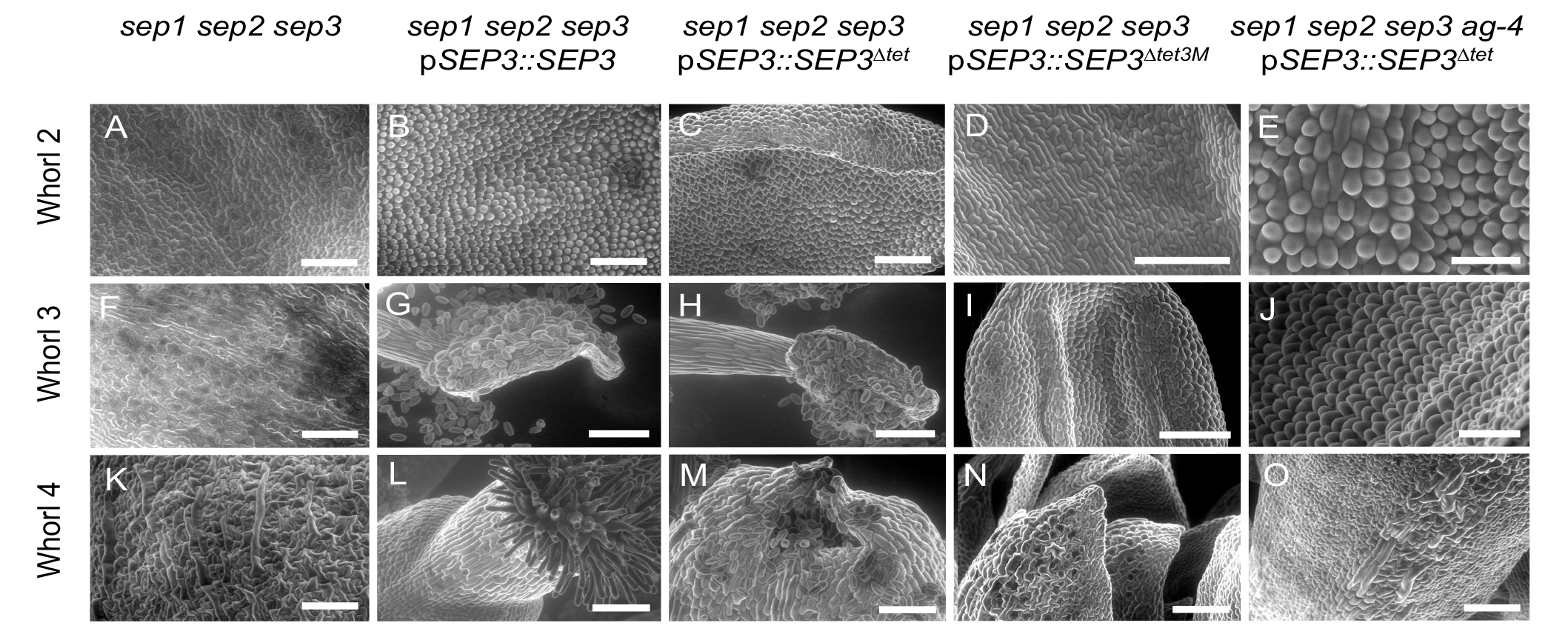
Scanning electron microscopy of epidermal cells for second, third and fourth whorl floral organs in *Arabidopsis* expressing wild-type and MADS mutants. **A-E**. SEM of adaxial cell surface of whorl 2, left to right, *sep1 sep2 sep3, sep1 sep2 sep3* expressing *SEP3, SEP3^Δtet^* or *SEP3^Δtet3M^*and *sep1 sep2 sep3 ag-4* expressing *SEP3^Δtet^* as labeled (top). Typical conical petal cells are observed in the *SEP3* (**B**) and *SEP3^Δtet^* (**C, E**) expressing lines, but absent in *SEP3^Δtet3M^* expressing lines (**D**). **F-J.** SEM of adaxial cell surface of whorl 3. Typical pollen grains are only observed in the *SEP3* (**G**) and *SEP3^Δtet^*(**H**) expressing lines. The triple mutant expressing *SEP3^Δtet3M^*(**I**) shows incomplete differentiation of the third whorl organs whereas the quadruple mutant expressing *SEP3^Δtet^* shows characteristic conical petal cells (**J**). **K-O.** SEM of the abaxial cell surface of whorl 4 in plants as in **A**. The triple mutant expressing *SEP3^Δtet3M^* (**N**) and the quadruple mutant expressing *SEP3^Δtet^* (**O**) exhibit elongated sepaloid cells and no stigmatic cells, whereas the triple mutant expressing *SEP3^Δtet^* exhibits partial complementation (**M**), with two unfused carpel with stigmatic cells present. Scale bars indicate 100 µm except for **E** (30 µm).

In order to further examine the role of tetramerization, the *sep1 sep2 sep3 ag-4* expressing *SEP3****^Δ^****^tet^* mutant was generated by crossing *sep1 sep2 sep3* expressing *SEP3****^Δ^****^tet^* and *sep1 sep2 ag-4*^+/-^. Due to the very low number of seeds produced, a single *sep1 sep2 sep3 ag-4* plant expressing *SEP3****^Δ^****^tet^* was genotyped and analyzed (Figures 4 and 5, fifth column). This mutant showed strong floral organ defects specifically in whorls 3 and 4, as would be expected due to both SEP3**^Δ^**^tet^ and AG**^Δ^**^tet^ exhibiting impaired tetramerization. In whorl 3, the stamens were replaced by six petaloid organs with conical cells characteristic of petals (Figures 4 and 5). As AG is required for repressing *AP1* expression in the third whorl, the lack of AG function due to impaired tetramerization would be predicted to result in petal formation instead of stamens in whorl 3, as shown in the *ag* loss-of-function mutants^38^. Whorl 4 was not complemented, showing an indeterminate flower consisting of sepaloid structures with characteristic elongated cells, as in *sep1 sep2 sep3* plants or *sep1 sep2 sep3* plants expressing *SEP3****^Δ^****^tet3M^* (Figures 4 and 5). Conversion of stamens to petaloid organs was also observed in 7 plants genotyped *sep1 sep2 sep3^+/-^ ag-4* expressing *SEP3****^Δ^****^tet^* (Supplemental Figure S3).

Taken together, these data demonstrate that perturbing MTF tetramerization by introducing structure-based mutations in SEP3 or in SEP3 and AG has a strong effect on floral organ differentiation and cell identity in the second, third and fourth whorls, correlating tetramerization defects characterized *in vitro* with physiological function.

## Discussion

MIKC^c^ MTFs fulfill important roles in plant reproductive development. Evidence from gymnosperms, angiosperms and ancestral reconstructions of the most recent common ancestor of extant seed plants suggests that tetramerization of MTFs is likely widespread, however whether or not tetramerization is required for specifying reproductive organ identity has been less clear^4, 10^. In mammals and fungi, for example, MTFs regulate different developmental processes via dimer formation, with no higher order MADS oligomerization states accessible or required for DNA-binding or activity^39, 40^. While the addition of the Keratin-like tetramerization domain occurred early in evolution, with MIKC^c^ MTFs even present in charophyte green algae, defining the physiological role of tetramerization has been challenging due to the difficulties in fully decoupling DNA binding and dimerization/tetramerization^11, 41^. In addition, *in vitro* studies of MADS tetramerization mutants have demonstrated robust DNA-binding of MADS homo-and heterodimers, further raising the question of whether or not tetramer formation is indispensable for physiological functions^30, 31, 42^.

In angiosperms, B and C class organ identity MADS are not able to tetramerize directly based on *in vitro* and *in vivo* studies, with tetramerization requiring a SEPALLATA clade member^42, 43^. Over-expression of A, B and C class MADS genes is not sufficient to confer organ identity, with conversion of leaves to petaloid or stamenoid organs requiring an E class MADS in addition to A, B and C class^22, 25^. However, SEP clade member also heterodimerize promiscuously with A, B and C MTFs, raising the possibility that SEP-containing MADS heterodimers are the essential complex for specifying organ identity^28^. Recent studies have further demonstrated that functional identity of MTFs is conferred at least in part by the dimerization I domain which helps determine MADS protein-protein interaction and DNA-binding specificity^12^. Combining structure-based mutagenesis, detailed *in vitro* characterization of oligomerization state, DNA-binding and comparative transgenic studies allows us to more fully determine the role of MADS tetramer formation in floral organ development. By progressively mutating the tetramerization interface and examining the DNA-binding patterns as well as the ability of *SEP3* mutants to complement the homeotic conversion of second, third and fourth whorl organs to sepals in the triple *sep1 sep2 sep3* mutant, the role of hetero-dimerisation versus heterotetramerisation of MADS organ identity complexes can be addressed. Based on the data presented here, second, third and fourth whorl organ identity requires tetramer formation of MTF complexes. While dimeric MADS complexes are able to strongly bind DNA *in vitro* and *in vivo* based on band shift assays, seq-DAP-seq and ChIP-seq experiments, this is not sufficient for proper gene regulation in the context of organ identity specification.

A key outstanding question is the underlying molecular mechanism of gene regulation by MADS tetrameric complexes. SEP3/AG wildtype complexes show an enrichment in 36, 46 and 56 base pair intersite spacing due to concurrent two-site binding of DNA by tetrameric complexes, with these distances present in genes important for meristem determinacy. Seq-DAP-seq studies demonstrate the loss of intersite spacing even for the weakly impaired SEP3**^Δ^**^tet^/AG tetramerization mutant, whose expression *in planta* led to an indeterminacy phenotype, correlating well with changes in regulation of genes such as *KNU* but no defects in second or third whorl organ specification and only limited defects in fourth whorl organ identity^30^. Importantly, however, examination of the genome-wide binding by the strong SEP3**^Δ^**^tet3M^/AG and SEP3**^Δ^**^tet3M^/AG**^Δ^**^tet^ tetramerization mutants in this study demonstrates the reduction in DNA-binding most strongly affects binding sites in regions enriched for specific intersite distances. These regions contain putatively relevant genes involved in organ development including *KAN2, JAG* and *INO*. Expression of strong SEP3 and AG tetramerization mutants *in planta* results in much more pronounced floral organ defects in addition to the indeterminacy phenotype observed for SEP3**^Δ^**^tet^-expressing plants. This may indicate that the experimental conditions of seq-DAP-seq are underestimating the ability of the SEP3**^Δ^**^tet^/AG complex to weakly tetramerize or that dimer binding, even to relatively poor binding sites that may require co-operativity *in vivo*, are detected in seq-DAP-seq, masking changes in binding at loci important for organ identity specification. In addition, an important limitation to seq-DAP-seq experiments is the use of naked DNA to examine binding patterns, thus neglecting the chromatin landscape, which plays a critical role in gene regulation. Recent studies have sought to address the challenge of deciphering the role of chromatin architecture in MTF gene regulation. *In vitro* and *in vivo* experiments for AP1 have shown that tetramerization of AP1 strengthens binding to CArG boxes on nucleosomal DNA and tetramer formation may be required for efficient displacement of histones for clustered MADS binding sites. Thus, optimized intersite spacing and nucleosome positioning may both be key to why tetramerization of MTFs is required *in vivo* for launching floral organ identity programs.

Taken together, the structural, *in vitro* and *in vivo* experiments presented here demonstrate the critical importance of MADS tetramer formation in floral organ identity in the second, third and fourth whorls, in addition to the previously described importance of tetramerization in floral meristem determinacy^30^. Interestingly, MIKC^c^ MTFs are present in non-seed plants including algae, mosses and ferns which implies that tetramerization may have occurred early in evolution and may be required for gene regulation for all MIKC^c^ MTFs in the green lineage, although this remains to be determined. Further studies examining the role of oligomerization and mechanisms of gene regulation in diverse species by MADS complexes will shed light on how this TF family has evolved central and diverse roles in development from algae to land plants.

## Materials and Methods

### Plant material and growth conditions

All experiments were performed using *Arabidopsis thaliana* WT and MADS mutants in the Col-0 background. The *ag-4* mutant, originally generated in the Ler background,^44^ was back-crossed 5 times in the Col-0 ecotype. The *ag*-*4* mutant expresses two variants of AG carrying deletion of 12 or 14 amino acids in the tetramerization interface, due to a splicing site mutation^44^. Seedlings were grown in controlled growth chambers in long day conditions (16h light**/**8h dark) at 22^◦^C for plant transformation and phenotype analysis.

### Plasmid construction for *sep1 sep2 sep3* and plant complementation analysis

The originally generated *sep1 sep2 sep3,* containing a T-DNA insertion in *SEP1* and an unstable transposon insertion in *SEP2* and *SEP3*^21^, was replaced in this study by a stable mutant generated using CRISPR-Cas9 genome editing to delete portions of the *SEP2* and *SEP3* genes. To generate a stable null mutation in *SEP2,* two guide RNA (gRNA) sequences were designed with no off targets using CHOPCHOP^45^. The two gRNA sequences were first cloned in pATU26:U26gRNA vectors and finally inserted into pCAMBIA together with the cassette containing the Cas9 sequence from pBSK:pUBQ10:CoCas9^46^ and transformed into *Agrobacterium tumefaciens*. The generated *sep2* mutant carries a deletion of 795 bp starting at +6 in exon 1 and removing the first 16 bp of exon 2, resulting in a frame shift. A similar strategy was followed to generate two *sep3* alleles. The first *sep3* mutant (*sep3-3*) carries a 1081 bp deletion removing the last 38 bp of intron 1 up to the first 83 bp of exon 8. The second generated s*ep3* mutant, named *sep3-4*, carries a deletion of 963 bp starting from +25 in exon 1. Sequences for generating gRNA are presented in Table SII. Sequences of *sep2* and *sep3* at the site of deletion are provided in Table SIII. The triple *sep1 sep2 sep3* mutants were generated by crosses. The newly generated mutants have the same flower phenotype of the previously described triple *sep1 sep2 sep3* transposon mutant, with sepaloid organs in all whorls and flower indeterminacy^21^.

For the complementation analysis, p*SEP3::SEP3* (ABRC stock number CD3-2708) and p*SEP3::SEP3^Δtet^* (ABRC stock number CD3-2709) were used. p*SEP3::SEP3^Δtet3M^* was constructed as described for the above plasmids using PCR amplified specific sequence of *SEP3^Δtet3M^* cloned into pSP64. These three plasmids allow the expression of *SEP3*, *SEP3^Δtet^* and *SEP3^Δtet3M^* under the control of the *SEP3* promoter and contain the *SEP3* regulatory intron 1 sequence cloned between exon 1 and 2, as described previously^30^. The vector backbone, pFP100, allows GFP expression in seeds for selection of transformants^47^.

### Plant transformation and floral phenotype analysis

For the generation of *sep1 sep2 sep3* expressing *SEP3*, *SEP3^Δtet^*and *SEP3^Δtet3M^,* heterozygous *sep1 sep2 sep3-3^+/−^* plants were transformed with the *pSEP3::SEP3*, *pSEP3::SEP3^Δtet^*, and p*SEP3:: SEP3^Δtet3M^*using the floral dip method^48^. Transformants were selected based on the fluorescence of GFP-positive seeds. For the generation of *sep1 sep2 sep3 ag-4* expressing *SEP3^Δtet^*, *sep1 sep2* was crossed with the *ag-4* mutant to generate the *sep1 sep2 ag-4*^+/-^ mutant. Pollen from *sep1 sep2 sep3-3* plants expressing *SEP3^Δtet^*was used to fertilize *sep1 sep2 ag-4^+/-^* and *sep1 sep2 sep3-3^+/-^ag-4^+/-^* plants expressing *SEP3^Δtet^*could be genotyped after crossing. Manual self-fertilization of these plants generated *sep1 sep2 sep3-3 ag*-*4* (named *sep1 sep2 sep3 ag-4* for simplicity) expressing *SEP3^Δtet^* in the next generation. All the primers used for plant genotyping are listed in Table SII.

Floral phenotypic analyses were performed by light microscopy on flower numbers 10–19 based on their order of emergence on T1 plants genotyped *sep1 sep2 sep3* expressing *SEP3* (3 T1), *SEP3^Δtet^* (2 T1) and *SEP3^Δtet3M^* (5 T1), on control untransformed *sep1 sep2 sep3* plants, and on *sep1 sep2 sep3 ag-4* expressing *SEP3^Δtet^* (1 line) and *sep1 sep2 sep3^+/-^ ag-4* expressing *SEP3^Δtet^* (7 lines). In Figure 4, black squares were added to mask magnification and scale marks automatically generated by the software and appropriate scale bars were added manually in white for clarity.

### Environmental scanning electron microscopy

Scanning electron microscopy (SEM) experiments were performed at the Electron Microscopy Facility of the Institut de Chimie Moleculaire of Grenoble Nanobio-Chemistry Platform, as previously described^12^. Untreated flowers were directly placed in the microscope chamber. Care was taken to maintain humidity during the pressure decrease in the chamber in order to prevent tissue drying. Secondary electron images were recorded with a Quanta FEG 250 (FEI) microscope while maintaining the tissue at 2 °C, under a pressure of 500 Pa and a 70% relative humidity. The accelerating voltage was 14 kV and the image magnification ranged from 100 to 800Å. Flowers from three independent lines were observed for each genotype.

### SEP3-AG K domain construct, protein expression and purification

The SEP3 K domain corresponding to residues 75-178 was PCR amplified and inserted by Gibson assembly to the NcoI/HindIII linearized pETDuet vector to generate the pETDuet-SEP3^75–178^ construct. A Tobacco Etch Virus (TEV) cleavable 6x histidine-maltose binding protein (His-MBP) tag amplified from the pETM-41 vector followed by the region corresponding to AG^90–189^ K domain with an additional TEV cleavage site at the C terminus, were inserted into the pETDuet -SEP3^75–178^ linearized by NdeI, using Gibson assembly to create the pETDuet SEP3^75–178^ /AG^90–189^ construct. Primers are listed in Table SII. *E. coli* BL21 Rosetta 2 (Novagen) were transformed with the pETDuet SEP3^75–178^ /AG^90–189^ construct and grown either in LB or minimal medium containing selenomethionine as described^49^. Cells were grown at 37 °C to an OD600 of 0.6**–**0.8 after which time the temperature was reduced to 18 °C and protein expression induced by addition of 1 mM of isopropyl-β-D-1- thiogalactoside for 12 h. Cells were harvested by centrifugation and the cell pellet resuspended in lysis buffer, 50 mM Tris-HCl pH 7.5, 300 mM NaCl, 1 mM tris(2- carboxyethyl)phosphine (TCEP), supplemented with 1x complete protease inhibitors (Roche).

Cells were lysed by sonication and cell debris pelleted at 25,000 rpm for 40 min. The soluble fraction was applied to a 1 ml Ni-NTA column, washed with lysis buffer + 10 mM imidazole and the protein eluted with lysis buffer + 250 mM imidazole. Cleavage of the His-MBP tag was carried out overnight at 4°C during dialysis against Tris-HCl 50 mM pH 7.5, 300 mM NaCl, 1 mM TCEP in the presence of 1:100 (w:w) His-tagged TEV protease. The protein was then passed over a Ni-NTA column to deplete the TEV and any uncleaved protein. SEP3^75–178^ /AG^90–189^ complex was further purified by gel filtration using a Superdex 200 10/300 column (GE Healthcare). The protein complex was concentrated to 6-8 mg/ml and used for crystallization trials.

### Protein crystallization, data collection and refinement

SEP3^75–178^ /AG^90–189^ at a concentration of 6-8 mg/ml was mixed at a 1:1 ratio with Tris-HCl 100 mM pH 8 and 2 M sodium formate. The protein crystallized after 3 days at 4 °C forming rectangle shaped single crystals. Seleno-methionine derivatized crystals were obtained after seeding with WT crystals. Glycerol was added to the drop to ∼20% final concentration as cryoprotectant and the crystals were then flash frozen in N_2(l)_. Diffraction data were collected at 100 K at the European Synchrotron Radiation Facility, Grenoble, France, on ID23-2 at a wavelength of 0.873 Å. Indexing was performed using MXCube^50^ and the default optimized oscillation range and collection parameters used for data collection. All datasets were integrated and scaled using the programs XDS and XSCALE^51^. For seleno-methionine containing crystals, 6 SeMet data sets were collected from three crystals. Data were automatically processed by XDS within the Grenades pipeline^52^ and submitted to CODGAS^53^ to group isomorphous datasets. This identified two datasets from the same crystal which were merged and analyzed by SIRAS using the CRANK2^54^ phasing program. Diffraction images and XDS input files have been deposited at Zenodo (). The partial model from CRANK2 was used for molecular replacement of the native dataset with Phaser^55^. Model building was performed using Coot^56^ and all refinements were carried out in Refmac^57^. The structure quality was assessed using MolProbity^58^. Data collection and refinement statistics are given in Table 1. The structure is deposited under PDB 8CRA.

### Plasmid construction and EMSA experiments

Vectors containing *AG* (At4g18960.1), *SEP3* (At1g24260.2), *SEP3^Δtet^*(At1g24260.3), *AP3* (At3g54340) and *PI* (At5g20240) cDNAs were used as previously described^30^. *AP1* (AT1G69120) cDNA was PCR-amplified using specific primers and inserted into XbaI/BamHI digested pSP64 (Promega) vector. Coding sequences for *SEP3^Δtet3M^* and *AG^Δtet^* were generated using the QuikChange (Agilent) protocol according to the manufacturer’s instructions and cloned into pSP64 vector as described for *AP1*. Primers used to generate the vectors are listed in Table SII. These vectors were used for *in vitro* protein production using SP6 High-Yield Wheat Germ Protein Expression System (Promega L3260) according to the manufacturer’s instructions. Electrophoretic mobility shift assay (EMSA) were performed as described^30^. The 103-bp DNA probe from the *SEP3* promoter^30^ containing two CArG box binding sites was labeled with Cy5 (Eurofins). For each EMSA, a negative control was run corresponding to labelled DNA incubated with *in vitro* transcription translation mix and empty pSP64 vector.

### Plasmid construction and seq-DAP-seq experiments

For seq-DAP-seq experiments, the following C-terminal-tagged constructs were generated using Gibson assembly and PCR amplified: pTnT*-SEP3^Δtet3M^-3FLAG* and pTnT*- AG^Δtet^- 5Myc* as described^31^. pTnT*-SEP3-3FLAG,* pTnT*-SEP3^Δtet^-3FLAG,* pTnT*-AG-5Myc* are reported previously^31^. Seq-DAP-seq for SEP3^Δtet3M^-AG complex and SEP3^Δtet^-AG^Δtet^ complex was performed as described previously^12, 31^. Briefly, 2 μg of each purified plasmid was used as input in a 50 μl TnT (Promega) reaction incubated at 25 °C for 2 h. The reaction solution was then combined with 50 μl IP buffer (PBS supplemented with 0.005% NP40 and proteinase inhibitors (Roche)) and mixed with 20 μl anti-FLAG magnetic beads (Merck Millipore M8823). Following 1 h incubation at room temperature, the anti-FLAG magnetic beads were immobilized, and washed three times with 100 μl IP buffer. TF complexes were eluted with 100 μl IP buffer supplemented with 200 μg/ml 3xFLAG peptide (Merck Millipore F4799). The eluted protein was then immobilized on anti-c-Myc magnetic beads (Thermo Fisher 88843) and washed three times with 100 μl IP buffer to isolate homogeneous SEP3^Δtet3M^-AG or SEP3^Δtet^-AG^Δtet^ complexes. The purified protein complexes, while still bound on anti-c-Myc magnetic beads, were incubated with 50 ng DAP-seq input library pre-ligated with Illumina adaptor sequences. The reaction was incubated for 90 min, and then washed six times using 100 μl IP buffer. The bound DNA was heated to 98 °C for 10 min and eluted in 30 μl EB buffer (10 mM Tris-Cl, pH 8.5). The eluted DNA fragments were PCR amplified using Illumina TruSeq primers for 20 cycles, and purified by AMPure XP beads (Beckman). The libraries were quantified by qPCR, pooled and sequenced on Illumina HiSeq (Genewiz) with specification of pairedend sequencing of 150 cycles. Each library obtained 10–20 million reads. The seq-DAP-seq was performed in triplicate.

### Seq-DAP-seq data analysis

For each seq-DAP-seq samples, reads were checked using FastQC^59^ and adaptor sequences removed with NGmerge^60^ and mapped with bowtie^61^ onto the TAIR10 version of the *A. thaliana* genome (https://www.arabidopsis.org), devoid of the mitochondrial and the chloroplast genomes. The duplicated reads were removed using the samtools rmdup program^62^.The resulting alignment files were used to derive the binding intensity of each complex at 6347 regions bound by the SEP3/AG complex^31^. The binding intensity of a given complex at bound regions was computed as the normalized reads coverage, averaged across replicates, and expressed as reads per kilobase per million mapped reads (RPKM). To limit the bias due to differences in the signal-to-noise ratio between seq-DAP-seq samples (Table SIV), the per-million scaling factor was done with the total number of reads mapped in peaks instead of all mapped reads. We made this choice over the classical normalization with all mapped reads because normalizing by total mapped reads flattens the signal for SEP3^Δtet^/AG and SEP3^Δtet3M^/AG (samples for these two conditions have the lowest fraction of reads in peaks (FRiP) values, Table SIII)^12, 63^. This artificially makes SEP3^Δtet^/AG^Δtet^ more similar to SEP3-AG. This choice assumes that differences in FRiP values are due to differential DAP-seq efficiency. The coverage fold reduction (CFR) was computed as the ratio between the mean normalized coverage of a complex relative to that of another complex. A SEP3/AG position weight matrix was used to search CArG boxes in the 6,367 bound sequences and subsequences with score > −9 were retained. This was used to separate regions harboring a preferred spacing from regions with no preferred spacing in figure 3B.

### Chip-seq experiments and data analysis

*sep1 sep2 sep3-4* lines expressing either wildtype *SEP3* or the tetramerization deficient, *SEP3^Δtet^* were used to conduct chromatin immunoprecipitation experiments according to previously published protocols^64^. Briefly, 1 g inflorescence (flower stage 1-12) were collected from 4–5-week-old plants. The tissue was fixed for 30 min and the immunoprecipitation performed using a SEP3-specific antibody followed by library preparation using ThruPLEX DNA-Seq Kit (Takara) and deep sequencing^65, 66^. Experiments were done with two biological replicates and the control sample was generated using pre-immune serum. The two lines were grown in parallel and genotyped (see primer Table SII) prior to sample collection. For each ChIP-seq data, reads were checked as described in the DAP-Seq data analysis section. Peaks were identified using MACS2^67^ and merged using MSPC^68^, resulting in 4,369 unique regions. The binding intensity of a given complex at bound regions was computed as the normalized reads coverage, averaged across replicates, and expressed as reads per kilo per million (RPKM).

### RNA-seq experiments and data analysis

Total RNA were extracted from two independent lines for *sep1 sep2 sep3* expressing *SEP3* and three independent lines for *sep1 sep2 sep3* expressing *SEP3^Δtet^*, and in duplicate from *sep1 sep2* and *sep1 sep2 sep3* lines, with lines as described ^30^. All the plants were grown in parallel. Quality of the total RNA was validated by their 260**/**280 absorbance ratio and the integrity of the ribosomal RNA by agarose gel. RNA libraries construction and sequencing were performed by GENEWIZ (USA) using Illumina HiSeq and 2 °ø 150bp configuration as described^31^. Between 25 and 35 million reads were obtained for each library. Mapping onto the Arabidopsis genome (TAIR10), read count per gene and statistical analysis were done using STAR (no multimapping, mismatch number < 10), FeatureCount (default parameters) and EdgeR (default parameters), respectively, available in the Galaxy platform^69, 70^. Genes were considered differentially expressed (DE) between two genotypes when the log FC was > 1 or <**−** 1 and the FDR value < 0.05. DE genes were determined for *sep1 sep2 sep3* expressing *SEP3^Δtet^* versus *sep1 sep2 sep3* and *sep1 sep2 sep3* expressing *SEP3* vs *sep1 sep2 sep3* expressing *SEP3^Δtet^*. DE genes were previously determined for *sep1 sep2 sep3* vs *sep1sep2* and *sep1 sep2 sep3* expressing *SEP3* vs *sep1 sep2 sep3*^31^.

### Data availability

Crystallographic data have been deposited with the PDB under the code 8CRA. RNA-seq, ChIP–seq and seq-Dap-Seq datasets have been deposited in the GEO database and can be download with the following tokens: ylszyocitlabxcx (RNA-seq), mrcricesndczrod (ChIP-seq) and clwncugebvmlrud (seq-DAP-seq).

### Funding

This project received support from the Agence National de la Recherche (ANR-16-CE92- 0023) and GRAL, a program from the Chemistry and Biology Health Graduate School of the University Grenoble Alpes (ANR-17-EURE-0003), with a thesis fellowship to AJ. The X-ray diffraction experiments were performed on beamline ID23-2 at the European Synchrotron Radiation Facility (ESRF), Grenoble, France. This work used the platforms of the Grenoble Instruct-ERIC center (ISBG; UAR 3518 CNRS-CEA-UGA-EMBL) within the Grenoble Partnership for Structural Biology (PSB), supported by FRISBI (ANR-10-INBS-0005-02). The research leading to these results has received funding from the European Community’s Seventh Framework Programme H2020 under iNEXT Discovery (project number 871037).

### Author contributions

V.H. and C.Z. conceived the study. V.H, C.Z and K.K. designed experiments. V.H., X.L., M.P., A.J., A.G., X.X., W.Y. performed the experiments. C.Z. and M.N. solved the 3D structure. R.B.-M, J.L., K.K. and F.P. analyzed the genome wide data. C.Z. and V.H. wrote the manuscript with the help of all authors.

## Acknowledgment

We thank Franck Wellmer (Smurfit Institute of Genetics, Dublin, Ireland) for providing the original *ag*-*4* mutant. The authors thank the NanoBio-ICMG Platform (UAR 2607, Grenoble) for granting access to the Electron Microscopy facility.

